# Phenology of plant reproduction, foliar infection, and herbivory change along an urbanization gradient

**DOI:** 10.1101/2022.03.22.485313

**Authors:** Quinn N. Fox, Mahal J. Bugay, Eleanor Grant, Olivia Shaw, Keiko Farah, Rachel M. Penczykowski

**Affiliations:** Department of Biology, Washington University in St. Louis, St. Louis, Missouri, 63130, USA

**Keywords:** fungal pathogen, global change ecology, human disturbance, plant development and life-history traits, plant phenology, plant-herbivore interactions, plant-pathogen interactions

## Abstract

- Urbanization involves numerous environmental changes that may affect the timing of plant reproduction and foliar damage by pathogens, herbivores, and human activities. Yet such relationships have not been examined simultaneously in plant populations across levels of urbanization.
- We conducted monthly surveys of 22 populations of *Plantago lanceolata* and *P. rugelii* in parks spanning an urbanization gradient. We quantified plant reproductive development and prevalence of powdery mildew infection, insect herbivory, and mowing damage. Additionally, we placed potted “sentinel” plants into field populations to directly measure infection and herbivory rates.
- Urbanization was associated with earlier flowering and more seed production for *P. rugelii*, but less seed maturation for *P. lanceolata*. Mildew epidemics on *P. rugelii* started earlier and achieved greater prevalence in more urban sites. Correspondingly, sentinels only became infected in suburban and urban sites. There was less infection on *P. lanceolata*, including sentinels, suggesting low availability of pathogen genotypes able to infect this species. Early-summer herbivory on both plant species was accelerated in urban sites.
- Urbanization has species-specific associations with reproductive phenology and is associated with increased early-summer herbivory, larger epidemics of a foliar pathogen, and more mowing damage on two weedy herbs

## Introduction

Life on Earth is becoming increasingly urbanized. As of 2018, 55% of the world’s population resides in urban areas, and that metric is anticipated to grow to 68% by 2050 (United Nations, 2018). Urbanization involves dramatic changes to the environment which can alter ecological and evolutionary processes and impact the fitness of organisms, including increases in habitat fragmentation, frequency of disturbance, coverage by impervious surfaces, temperature, pollution, and introduction of invasive species (Johnson & Munshi-South, 2017). A meta-analysis of >100 published studies revealed wildlife face significantly greater risk of parasite infection in urban areas (Murray et al., 2019). However, effects of urbanization on plant fitness are poorly understood (Johnson & Munshi-South, 2017; Rivkin et al., 2019). Only one study has tested effects of urbanization on tree diseases (van Dijk et al., 2022), and none have tested effects of urbanization on diseases of wild herbs. No studies have simultaneously quantified the relationships between urbanization and the timing of plant reproduction and damage from pathogens, herbivores, and human activities.

A well-known consequence of urbanization is the urban heat island effect (UHI), in which cities are warmer than surrounding environments due to retention of solar radiation by urban materials and release of heat from high densities of vehicles and other machinery (Zhou et al., 2017). These increased temperatures can impact plant development; in a study of herbarium records, 24% of herbaceous and woody plant species flowered earlier in urban environments (Neil et al., 2010). Whether UHI advances plant phenology depends on baseline climate in the region; flowering date advances with city size only in regions with cool or cold winters, where degree-days are limiting to development (Li et al., 2019).

Urban heating may increase or decrease prevalence of plant diseases, depending on regional climate, magnitude of UHI, and thermal sensitivity of host and pathogen physiological traits. Here, we focus on fungal pathogens because these are the most well studied in wild plant populations (Alexander, 2010; Burdon & Laine, 2019). Across life history processes, plant-associated fungi typically have temperature minima of 5-15 °C, optima of 18-28 °C, and maxima of 26-38 °C (Chaloner et al., 2020). In regions where baseline climate is cooler than optimal for a pathogen, UHI may promote disease. Indeed, UHI is linked to increased severity of powdery mildew on English oak in Europe (van Dijk et al., 2022). However, in regions that are already warmer than optimal for a pathogen, additional heat in cities may inhibit pathogen growth. Effects of UHI on disease can further depend on microclimate heterogeneity (Penczykowski et al. 2018), temporal climatic variation (Egerer et al., 2020), and thermal sensitivity of plant defense responses (Desaint et al., 2021; Wang et al., 2009).

Fragmentation of urban green spaces can lead to greater isolation of plant populations compared to more continuous rural habitats (Dubois & Cheptou, 2017). Alternatively, plant or pathogen species that grow or disperse along roadways may be more highly connected in cities with denser road networks. Fungal infection is more likely in populations of ribwort plantain along roadsides, particularly at hubs of a road network, suggesting that vehicular traffic facilitates spore transport (Numminen & Laine, 2020). Increasing road density and traffic volume from rural to urban environments could therefore drive increases in fungal infection prevalence.

Rates of insect herbivory can also be altered by abiotic (e.g., temperature, light, and sound) and biotic (e.g., biodiversity of plants, insects, and predators) features of cities (Miles et al., 2019). UHI may accelerate emergence of insect herbivores in spring (Bale et al., 2002). If host plant phenology is similarly accelerated, this should lead to higher rates of herbivory earlier in the growing season. Contrastingly, if UHI causes phenological asynchrony between insects and their host plants less herbivory is expected (Bale et al., 2002). From the few published studies linking urbanization to herbivory both positive (Cuevas-Reyes et al., 2013; Just et al., 2019; Moreira et al., 2019) and negative (Meineke et al., 2019) relationships been documented (Miles et al., 2019). Other evidence for changes in herbivory pressure comes from declines in production of an antiherbivore chemical defense (hydrogen cyanide) in white clover with urbanization (Johnson et al., 2018; Santangelo et al., 2022).

The frequency and intensity of disturbance from human activities may also increase with urbanization. For herbs growing along roadways or lawns, human activities may damage or remove leaves and reproductive tissues directly or affect the probability of damage by pathogens or herbivores. On golf courses, increased human footsteps and lowered mower blade height increased severity of fungal infections on grass (Inguagiato et al., 2009; Roberts & Murphy, 2014; Williams et al., 2001). Effects of walking and mowing on plant populations across levels of urbanization remain to be explored.

We examined the phenology of plant reproduction, fungal infection, insect herbivory, and lawn mowing damage across a 56-km land use gradient in the St. Louis metropolitan area in Missouri, United States (Fig. **1**, Table **S1**). We conducted monthly field surveys of *Plantago lanceolata* (ribwort plantain) and *P. rugelii* (blackseed plantain) populations co-occurring in 22 sites in summer–fall 2019 and 2020. In a field experiment performed in 2020, we placed groups of healthy, pathogen-free “sentinel plants” of these species into a subset of sites to directly measure rates of infection and herbivory across the urbanization gradient. We hypothesized that urbanization would be associated with: (1) earlier flowering and seed production, (2) earlier and larger powdery mildew epidemics, (3) earlier and overall more prevalent insect herbivory, and (4) more mowing damage on leaves. We further hypothesized that rates of infection and herbivory on sentinels would positively correlate with prevalence of infection and herbivory in the field populations where they were placed.

**Fig. 1.**
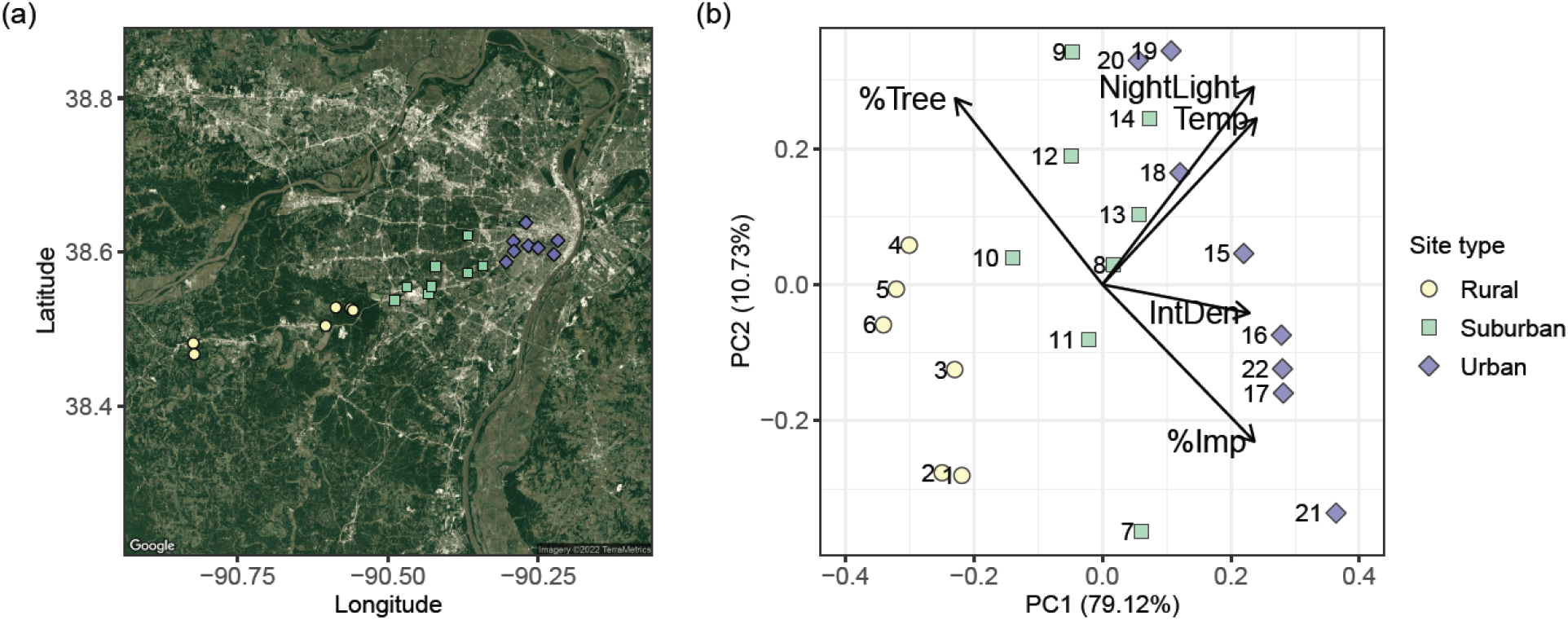
(**a**) Google Earth satellite view of the focal urbanization gradient in St. Louis, Missouri, USA. Urban sites are plotted as purple diamonds, suburban as green squares, and rural as yellow circles. (**b**) Biplot of a principal component analysis of environmental variables at the study sites. Black arrows represent loadings of percent impervious area (%Imp), percent tree cover (%Tree), estimated intersection density of walkable roads (IntDen), average temperature from June–October (Temp), and nighttime radiance (NightLight). Sites are numbered from west to east (see Table **S1** for additional site information).

## Materials and Methods

### Study system

This study focused on two co-occurring herbs, *Plantago lanceolata* (ribwort plantain) and *Plantago rugelii* (blackseed plantain). These short-lived perennials grow as rosettes in pastures or human-disturbed landscapes (e.g., lawns and roadsides), and are tractable model organisms for studies of plant–pathogen and plant–herbivore interactions across land use gradients (Penczykowski & Sieg, 2021). *Plantago lanceolata* is native to Eurasia but cosmopolitan in its distribution, and *P. rugelii* is endemic to eastern North America (Penczykowski & Sieg, 2021).

The most globally and locally common foliar fungal pathogens of *Plantago* are the specialist powdery mildews *Podosphaera plantaginis* (Castagne; U. Braun and S. Takamatsu) and *Golovinomyces sordidus* (L. Junell) V.P. Heluta (Braun & Cook, 2012). Powdery mildews – obligate fungal pathogens in the order Erysiphales – grow on the surface of leaves and extract nutrients from their host’s epidermal tissue (Bushnell, 2002). Chains of asexual spores produced from mycelia on the leaf surface give infected leaves a white, dusty appearance. The spores are passively transmitted via wind, and more than 90% land within 2 m of the source plant (Tack et al., 2014). However, occasional long-distance spore transport allows for pathogen persistence at the regional scale (Ovaskainen & Laine, 2006). Powdery mildews overwinter via sexual resting structures that release spores when conditions are favorable in spring (Tack & Laine, 2014).

*Plantago* are also host to numerous invertebrate herbivores (Penczykowski & Sieg, 2021). This study focuses on leaf mines and chewing damage. Leaf mines on *Plantago* are created by larvae of the leaf miner fly *Phytomyza plantaginis* (Penczykowski & Sieg, 2021). Chewing damage on leaf edges or between leaf veins could be due to a variety of arthropod taxa, particularly larval lepidopterans. In the region of our study, these include specialist common buckeye caterpillars (*Junonia coenia*) as well as generalist species (Penczykowski & Sieg, 2021).

### Focal populations

We surveyed 22 sites (parks and nature reserves) with co-occurring *P. lanceolata* and *P. rugelii* populations extending southwest from downtown St. Louis City along the I-44 interstate highway corridor (Fig. **1a**, Table **S1**). In a few large parks or nature areas, we surveyed two sites separated by at least 0.45 km. The study began with 19 sites in July 2019, two sites were added at Shaw Nature Reserve in August and September 2019, and a site was added at Forest Park in June 2020 (Table **S1**).

We classified eight sites within St. Louis City as “urban”, eight sites in St. Louis County east of Missouri Route 141 as “suburban”, and six sites west of Missouri Route 141 as “rural”. These classifications were based on a principal component analysis (PCA) of environmental and spatial variables performed with the ‘prcomp’ function in R version 3.6.2 (R Core Team, 2019). The PCA included average temperature from June through October (Temp), nighttime radiance (NightLight), percent impervious area (%Imp), percent tree cover (%Tree), and estimated intersection density of walkable roads (IntDen; walkable roads defined as having speed limits between 8-55 miles per hour) (Fig. **1b**). The temperature data represented long-term (1970-2000) averages at these sites during the focal months of our study (WorldClim 2; Fick & Hijmans, 2017). Nighttime radiance data came from the NASA Black Marble data product for December 2020 (NASA Worldview).

For most sites, percent impervious area, percent tree cover, and intersection density were obtained through the EnviroAtlas Community dataset for St. Louis (US EPA, 2021a,b,c). Some rural sites were located outside the boundaries of that urban-centered dataset. Thus, for rural sites #1, 2, and 3 (Table **S1**), we estimated percent impervious area and percent tree cover from the National Land Cover Database (MRLC, 2019). Specifically, we used the Summarize Categorical Raster tool in ArcGIS Pro to extract land use pixel count data from the NLCD layer for the census blocks containing the sites. We calculated percent impervious area as percent of pixels in low, medium, or high intensity development classes. Similarly, we calculated percent tree cover as the percent of pixels in deciduous forest, evergreen forest, mixed forest, shrub scrub, or woody wetlands classes. Intersection density data was not available through EnviroAtlas for rural sites #1, 2, 3, 5, or 6 (Table **S1**) thus we estimated intersection density using the streets layer from the ESRI Streetmap Premium dataset as a base (ESRI, 2020). After limiting to streets with speed limits between 6-55 mph, we manually digitized all intersections within a 750-m radius of the focal sites, and used the Kernel Density tool to generate an intersection density layer. From that layer, we extracted values of intersection density at the study sites, following methods in the EnviroAtlas metadata (US EPA, 2021c).

### Field survey methods

We performed monthly surveys in the focal populations from July to October 2019 (four surveys) and from June to October/November 2020 (five surveys). This time frame was chosen to represent the majority of the growing season of the plants, their fungal pathogens, and their insect herbivores. In each survey, we collected data for 50 individuals each of *P. lanceolata* and *P. rugelii*. On a meandering path through each population, we selected the nearest plant of either species that was at least 1 m from the previously surveyed individual of that same species. The average sizes of the surveyed areas were 1004, 1714, and 2699 m^2^ for rural, suburban, and urban sites, respectively (Fig. **S1**; sizes estimated from polygons in Google Earth). Rural survey areas tended to be smaller because those sites were typically bounded by tall grassland or woodlands where *Plantago* do not grow. By contrast, urban and suburban sites typically had larger total areas with *Plantago* present, and our surveys covered more ground.

For each selected plant, we recorded the status of the most mature flower stem on the plant, if present, using the following ordered categories: (0) stem present but flowers/seeds removed by mowing, (1) budding, (2) flowering, (3) immature seeds, (4) mature seeds, or (5) seeds dispersed. Seeds were categorized as immature if the seed capsules were green and plump, and mature if they were brown and papery. We recorded the powdery mildew infection status for each plant as the presence or absence of conspicuous white mycelia and/or conidia on the leaf surface. In 2020, to assess whether the infection statuses of the selected plants were representative of nearby conspecifics, we additionally recorded the numbers of infected conspecifics within a 1.5-m radius of each focal individual, using the following categories: “0” = 0, “1” = 1-10, “2” = 11-50, “3” = 51-100, and “4” = 101 or more plants. In 2020, we recorded the presence or absence of two main types of herbivory damage (chewing and leaf mines on leaves) and mowing damage (straight cuts across leaves).

### Sentinel experiment

In July 2020, healthy, greenhouse-grown “sentinel” plants were placed into 15 field survey sites to directly test rates of powdery mildew infection and herbivory across the urbanization gradient. These sites (five of each site type) were chosen from the full set of 22 sites based on presence of mildew on at least one of the *Plantago* species in 2019. We selected sites based on that criterion to avoid biasing the outcome of the experiment towards rejection of the null hypothesis that infection prevalence would be the same across site types.

Sentinel plants were grown from *P. lanceolata* and *P. rugelii* seeds collected from wild, uninfected plants in St. Louis City and County during July–October 2019. To capture variation in plant genetic background, we planted seeds from six maternal lines of *P. lanceolata*, and five of *P. rugelii*. Each maternal line consisted of a single seed spike from a mother plant. Seeds were sown in BM6 All-Purpose soil (Berger) on 16 May 2020 in the Jeanette Goldfarb Plant Growth Facility at Washington University in St. Louis. All six *P. lanceolata* lines but only three of the *P. rugelii* lines successfully germinated. On 1 June, we moved the seedlings to a hoop house at Tyson Research Center (Eureka, Missouri) and transplanted each into a 4.5-in diameter pot. To ensure that sentinels remained uninfected before deployment into the field, we covered each with an autoclaved, spore-proof pollination bag (PBS International, model 3D.55), which we secured around the pot with a silicone band. Three times per week, we placed the sentinels in a shallow pool of water for 10 min, allowing water to soak up through the base of the pot.

Sentinels were deployed into field sites between 7-9 July 2020. Due to much higher germination success for *P. lanceolata* than *P. rugelii*, we were able to place 20-23 *P. lanceolata* into each of 15 sites (n = 322) but only 3-4 *P. rugelii* into each of 9 sites (n = 36). Within sites, sentinels were randomly divided between three seedling trays at the base of a single tree (i.e., all sentinels were at one shaded location per site). We removed the pollination bags and collected initial data on the number of leaves, length and width of largest leaf, and flower maturity stage. We confirmed that no sentinels were infected with powdery mildew, and recorded any existing damage on leaves (e.g., from greenhouse or hoop house insects encountered prior to enclosure with pollination bags). We also recorded the distance from the trays of sentinels to the nearest infected *Plantago* in the surrounding wild population. We placed a temperature datalogger (HOBO MX2201) just below the soil surface in one pot per tray to monitor temperature differences between sites. The trays were filled with an inch of water at the time of deployment and once the following week. It did not rain during the week of the experiment, so this watering scheme achieved healthy moisture levels in the pots.

Between 14-16 July 2020, sentinels were retrieved from the field sites in the same order they were deployed. Before removal from the field, we recorded the number of leaves, length and width of largest leaf, and presence/absence of chewing and leaf miner herbivory for each leaf. Then we re-covered each sentinel with a pollination bag and moved them to a common location 5 km from the nearest field site to allow any powdery mildew infections to develop for another week. During that week, we watered plants twice (in shallow pools for 10 min), and at the end of the week we recorded infection status of each leaf.

### Statistical analyses

All statistics were performed in R version 3.6.2 (R Core Team, 2019). Field survey data were analyzed separately for each species and year. In models testing effects of survey month (ordered factor), site type (urban, suburban, or rural), and their interaction on a given response variable, we included site (population) identity as a random effect. We performed post-hoc Tukey’s tests of contrasts between site types within each month if there was at least a marginally significant (P < 0.10) site type x month interaction, or between site types averaged across months if there was a significant (P < 0.05) main effect of site type but no interaction (package ‘emmeans’; Lenth, 2022). We report all site type contrasts with P < 0.10, reserving the word “significant” for P < 0.05.

We analyzed plant reproductive development stage as an ordered categorical response variable (levels 0–5, as explained above in *Field survey methods*) in cumulative link mixed models (package ‘ordinal’; Christensen, 2019). Prevalences of powdery mildew infection, leaf mines, and mowing were logit-transformed and analyzed in linear mixed effects models (package ‘nlme’; Pinheiro et al., 2012) with first order autocorrelation structure. For prevalence values of zero or one, we added (to zeros) or subtracted (from ones) 0.0001 before logit-transforming. We detected no temporal autocorrelation in prevalence of chewing herbivory, so this was analyzed in a generalized linear mixed model (GLMM, package ‘lme4’; Bates et al., 2015) with binomial error distribution and logit link function.

To test if the infection statuses of the surveyed plants were representative of nearby conspecifics, we used binomial generalized linear models (logit link). Specifically, we modelled binary infection status of surveyed plants in response to the categorical abundance of infected conspecifics in the surrounding 1.5 m radius, with site identity as a random effect.

In the sentinel experiment, we analyzed the proportion of leaves on each plant with powdery mildew, leaf mines, and chewing damage (i.e., “severity” of each damage type) using binomial GLMMs (logit link) with random effects of tray identity nested within site identity (because there were three trays of sentinels in each field site). In addition to testing effects of site type on infection and herbivory severity, we included covariates reflecting the initial size of sentinels. Specifically, these were scores on the first two principal components axes from a PCA of number of leaves and length and width of longest leaf at the start of the experiment. For *P. lanceolata*, the first principal component axis (PC1) explained 65.06% of variation in initial plant size data and the second (PC2) explained 34.72%; for *P. rugelii*, PC1 explained 75.36% and PC2 explained 21.92% of variation. For analysis of infection severity, we included distance to nearest infected wild plant as a covariate. For analyses of severity of leaf mines and chewing, we additionally included the initial severity (chewing) or presence/absence (mines) of these damage types as covariates, to account for any herbivory that occurred prior to placement in the field. For both infection and herbivory, we tested whether prevalence on sentinels was correlated with prevalence in the wild populations where they had been placed.

## Results

### Field site characteristics

In the PCA of environmental variables at the study sites, the first two principal components (PC1 and PC2) together explained 89.85% of the total variation in the data (Fig. **1b**). The pre-classified site types (rural, suburban, and urban) were primarily separated along PC1, which was associated with greater temperature, nighttime radiance, intersection density, and percent impervious area, and with lower percent tree cover (Fig. **1b**). Variation within site types occurred primarily along PC2, which was associated with greater temperature, nighttime radiance, and percent tree cover, and with lower percent impervious area (Fig. **1b**).

### Field surveys: plant reproductive phenology

In both 2019 and 2020, *P. lanceolata* populations had begun flowering in all site types before our first survey, the frequency of flowering peaked in July, and frequency of mature and/or dispersed seeds peaked in August (Fig. **2a**,**c**). For *P. lanceolata* in 2019, parameters could not be uniquely determined for the model of reproductive development through time when October data were included. Excluding October 2019 from the analysis, the effect of site type on *P. lanceolata* reproductive development (as an ordered categorical variable) varied through time in both years (site type x month interaction in 2019: Likelihood Ratio [LR] □^2^ = 20.40, df = 4, P = 0.0004; in 2020: LR □^2^ = 51.52, df = 8, P < 0.0001). Post-hoc Tukey’s tests showed that, in 2019, maturity was marginally lower in urban and/or suburban than rural sites in July (urban-rural: P = 0.050, suburban-rural: P = 0.068) and August (suburban-rural: P = 0.060), with no other differences between site types within a given month (P > 0.10 for all unreported pairwise contrasts; Fig. **2a**). In 2020, *P. lanceolata* reproductive development was marginally lower in urban than rural sites in August (P = 0.079) and significantly lower in urban than either suburban or rural sites in September (urban-rural: P = 0.036, urban-suburban: P = 0.008; Fig. **2c**).

**Fig. 2.**
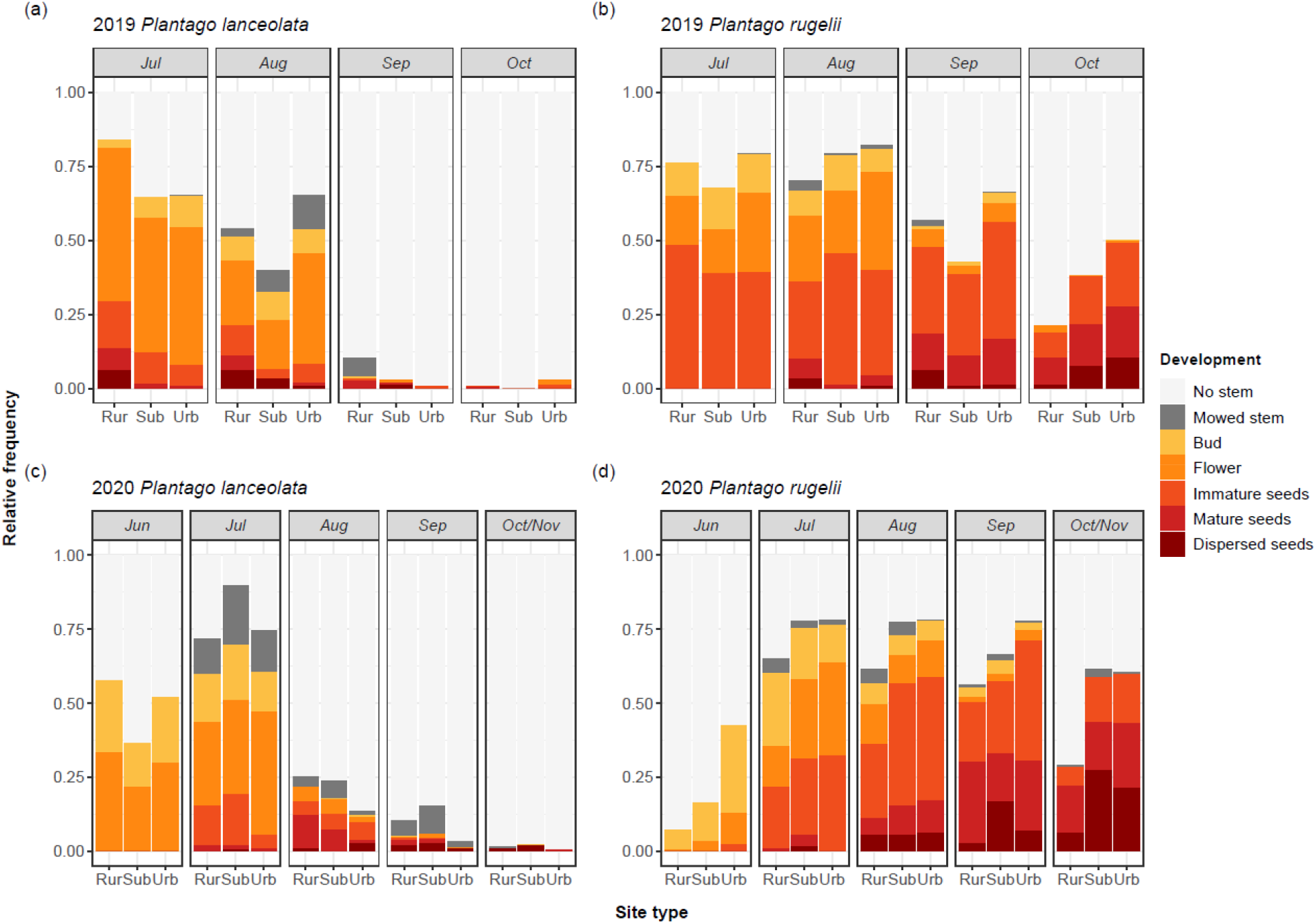
Relative frequency of each stage of plant reproductive development for each site type and survey month in 2019 and 2020 (Rur = rural, Sub = suburban, Urb = urban).

In both years, flowering continued at a higher rate into August for *P. rugelii* compared to *P. lanceolata*, and overall rates of seed production and maturation were also much higher for *P. rugelii* (Fig. **2**). The effect of site type on *P. rugelii* reproductive development varied through time in both years (site type x month interaction in 2019: LR □^2^ = 115.72, df = 6, P < 0.0001; in 2020: LR □^2^ = 128.98, df = 8, P < 0.0001; Fig. **2b**,**d**). Post-hoc tests revealed that *P. rugelii* development was significantly lower in suburban than either rural (P = 0.0092) or urban (P = 0.0002) sites in September 2019 but increased with level of urbanization in October 2019 (suburban-rural: P = 0.0001, urban-rural: P < 0.0001, urban-suburban: P = 0.009; Fig. **2b**). Our June 2020 survey captured significantly earlier *P. rugelii* development with increasing levels of urbanization (suburban-rural: P = 0.018, urban-rural: P < 0.0001, urban-suburban: P = 0.0011; Fig. **2d**). Maturity of *P. rugelii* was greater in urban than rural sites in both August and September 2020 (P = 0.025 and P = 0.043, respectively), and greater in both suburban and urban than rural sites in October/November 2020 (P < 0.0001 for both contrasts).

### Field surveys: powdery mildew infection

There was very little powdery mildew infection on *P. lanceolata* in 2019, and generally small epidemics in 2020 (Fig. 3a,c). For this species, there were no significant site type x month effects on infection prevalence (in 2019: LR □^2^ = 4.00, df = 6, P = 0.68; in 2020: LR □^2^ = 10.34, df = 8, P = 0.24), nor significant main effects of site type (in 2019: LR □^2^ = 2.43, df = 2, P = 0.30; in 2020: LR □^2^ = 3.36, df = 2, P = 0.19). There was a significant main effect of month in 2020 only (in 2019: LR □^2^ = 3.92, df = 3, P = 0.27; in 2020: LR □^2^ = 47.64, df = 4, P < 0.0001).

**Fig. 3.**
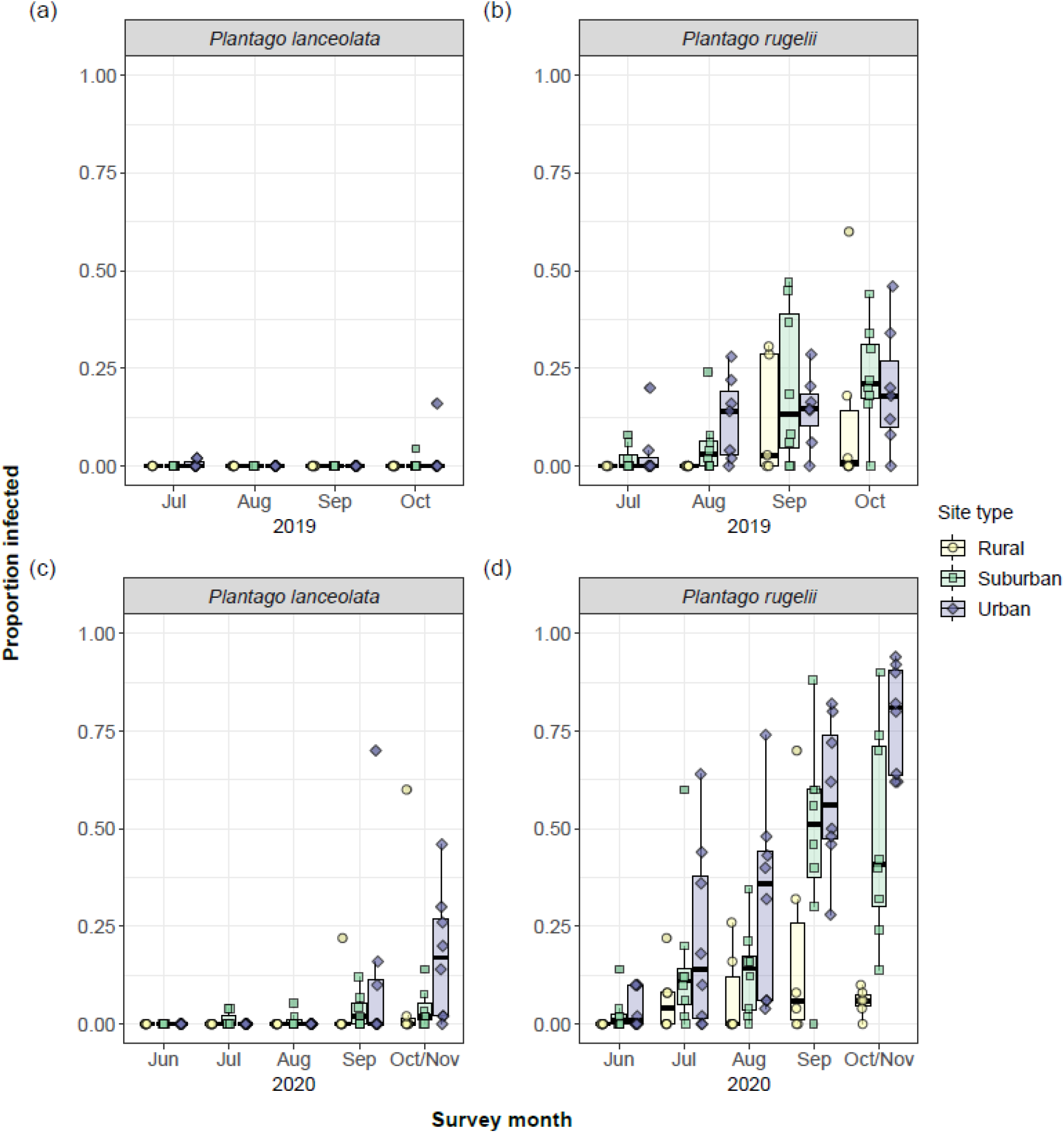
Proportion of plants infected with powdery mildew in each population. Box-and- whisker plots are grouped by site type and survey month in 2019 and 2020.

There were large powdery mildew epidemics on *P. rugelii* in both years, and epidemics started earlier in suburban and urban populations (Fig. **3b**,**d**). In both years, *P. rugelii* infection prevalence was significantly affected by site type (in 2019: LR □^2^ = 8.89, df = 2, P = 0.012; in 2020: LR □^2^ = 22.00, df = 4, P < 0.0001) and month (in 2019: LR □^2^ = 22.57, df = 3, P < 0.001; in 2020: LR □^2^ = 70.42, df = 4, P < 0.001), but there were no site type x month interactions (in 2019: LR □^2^ = 4.39, df = 6, P = 0.62; in 2020: LR □^2^ = 3.90, df = 6, P = 0.87). For 2019, post-hoc testing revealed significantly greater infection prevalence in urban than rural sites (suburban-rural: P = 0.067, urban-rural: P = 0.037; Fig. **3b**). In 2020, there was significantly greater infection prevalence in both suburban and urban relative to rural sites (suburban-rural: P = 0.0061, urban-rural: P = 0.0007; Fig. **3d**).

The infection status of surveyed plant individuals was consistent with that of nearby plants in the population. Specifically, presence/absence of powdery mildew infection on surveyed plants was significantly positively related to the abundance of infected conspecifics within a 1.5-m radius (*P. lanceolata:* □^2^ = 262.81, df = 3, P < 0.0001; *P. rugelii:* LR □^2^ = 1097.6, df = 3, P < 0.0001; Fig. **S2**).

### Field surveys: herbivory

For both species, prevalence of leaf mines depended on the interaction of site type and month (*P. lanceolata*: □^2^ = 16.37, df = 8, P = 0.037; *P. rugelii*: □^2^ = 55.84, df = 8, P < 0.0001; Fig. **4a**,**b**), and the only significant differences between site types occurred in June. For *P. lanceolata*, there was significantly greater prevalence of leaf mines in urban than suburban sites in June (urban-rural: P = 0.092, urban-suburban: P = 0.0083). For *P. rugelii*, there was greater prevalence of leaf mines in urban than either suburban or rural sites in June (urban-rural: P = 0.0017, urban-suburban: P = 0.0001).

**Fig. 4.**
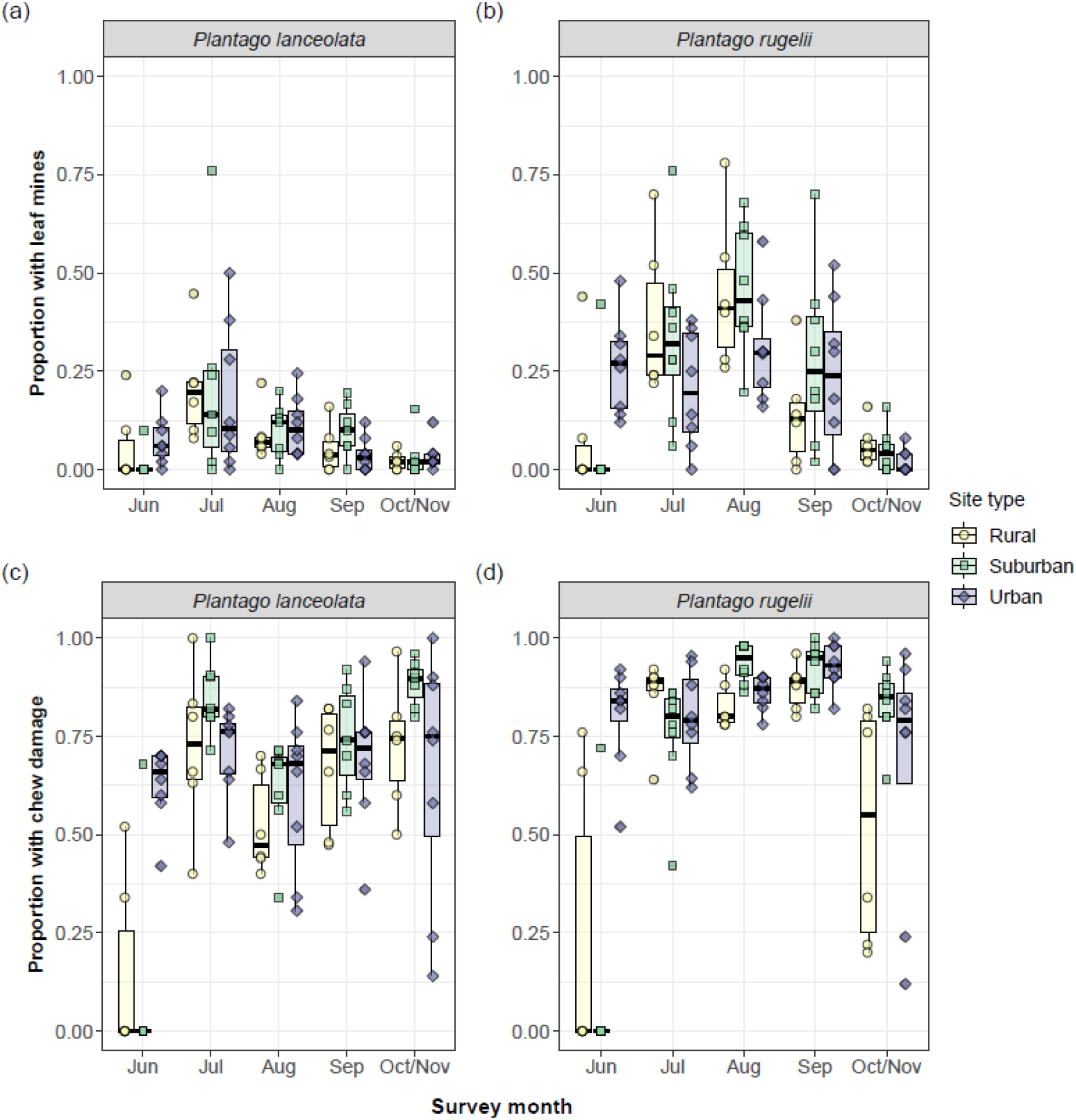
Proportion of plants with chewing damage or leaf mines in each population. Box- and-whisker plots are grouped by site type and survey month in 2020.

Chewing-type herbivory also depended on the interaction between site type and month (*P. lanceolata*: □^2^ = 66.17, df = 8, P < 0.0001; *P. rugelii*: □^2^ =88.51, df = 8, P < 0.0001; Fig. **4c**,**d**). For both species, there was greater prevalence of chewing damage in urban than suburban or rural sites in June (*P. lanceolata*: urban-rural: P < 0.0001, urban-suburban: P < 0.0001; *P. rugelii*: urban-rural: P < 0.0001, urban-suburban: P < 0.0001).

### Field surveys: mowing

For *P. lanceolata*, prevalence of mowing damage on leaves depended on site type (□^2^ = 7.50, df = 2, P = 0.024) and month (□^2^ = 43.10, df = 4, P < 0.0001), but not their interaction (□^2^ = 10.28, df = 8, P = 0.25). For this species, there was marginally greater mowing prevalence in suburban and urban than rural sites (suburban-rural: P = 0.049, urban-rural: P = 0.083; Fig. **5a**).

**Fig. 5.**
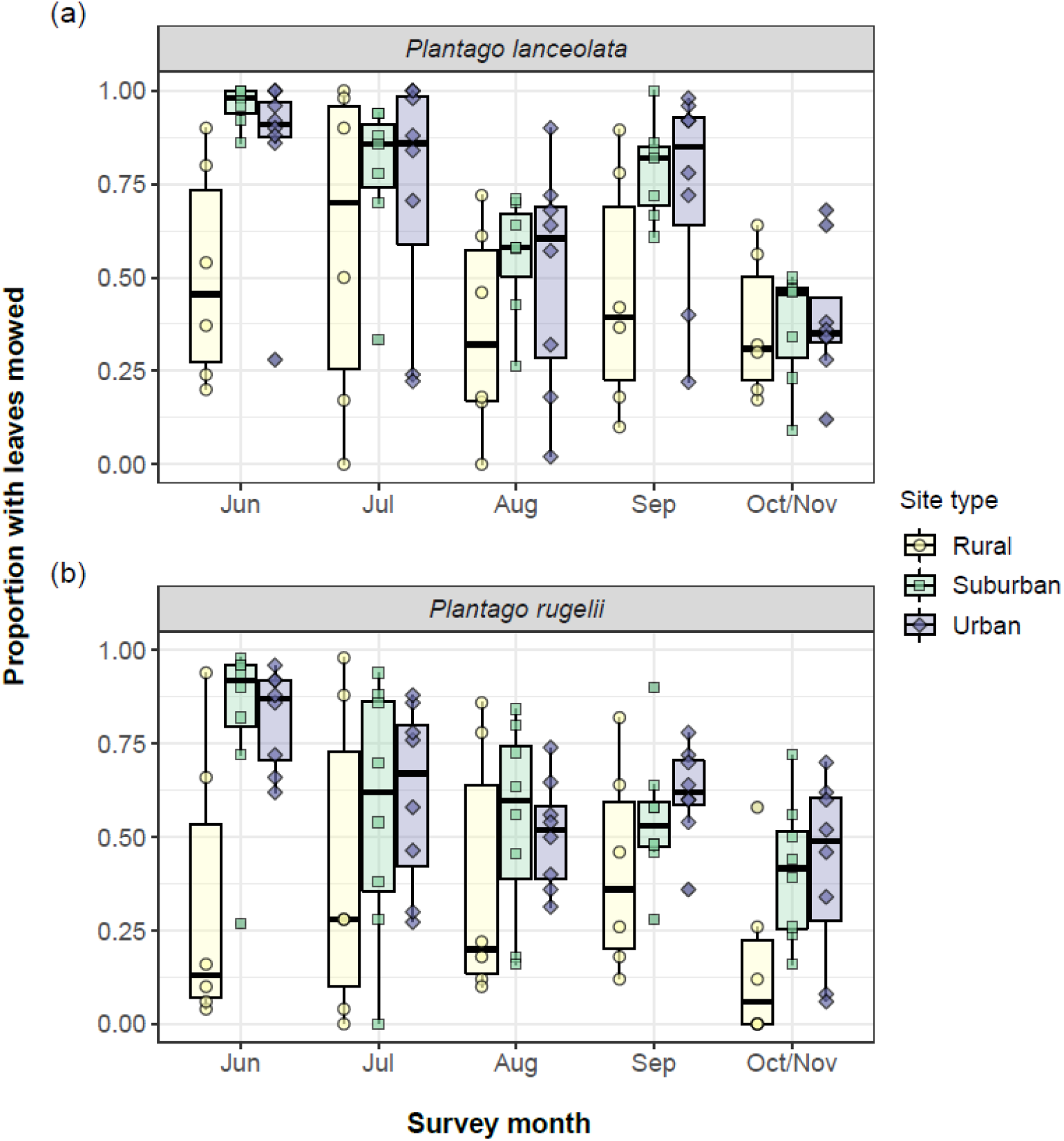
Proportion of plants with leaf damage from lawn mower blades in each population. Box-and-whisker plots are grouped by site type and survey month in 2020.

For *P. rugelii*, there was a marginally significant site type x month effect on prevalence of mowing damage (site type x month: □^2^ = 13.79, df = 8, P = 0.087; site type: □^2^ = 8.56, df = 2, P = 0.014; month: □^2^ = 23.44, df = 4, P = 0.0001; Fig. **5b**). This weak interactive effect reflected marginally greater mowing prevalence in suburban and urban relative to rural sites in June (suburban-rural: P = 0.034, urban-rural: P = 0.062) and significantly greater mowing prevalence in suburban and urban relative to rural sites in the final October/November survey (suburban-rural: P = 0.0020, urban-rural: P = 0.0025; Fig. **5b**).

### Sentinel experiment: powdery mildew infection and herbivory

All 322 *P. lanceolata* sentinels survived to the end of the experiment, and only one became infected with powdery mildew (Fig. **6a**). This infected sentinel was placed in Buder South Park within 1.5 m of the only wild infected *P. lanceolata* individual we found after a thorough search at that site (Fig. **S3**; note that the one wild infected plant was not among the 50 plants surveyed at that site in July 2020, so infection prevalence of wild plants was recorded as zero for that site-date; Fig. **6a**). No *P. lanceolata* sentinels became infected in Tilles Park, which did have a small number of infected wild plants at that time (Fig. **6a**), likely because the sentinels were located almost 6 m from the nearest wild infected plant (Fig. **S3**).

**Fig. 6.**
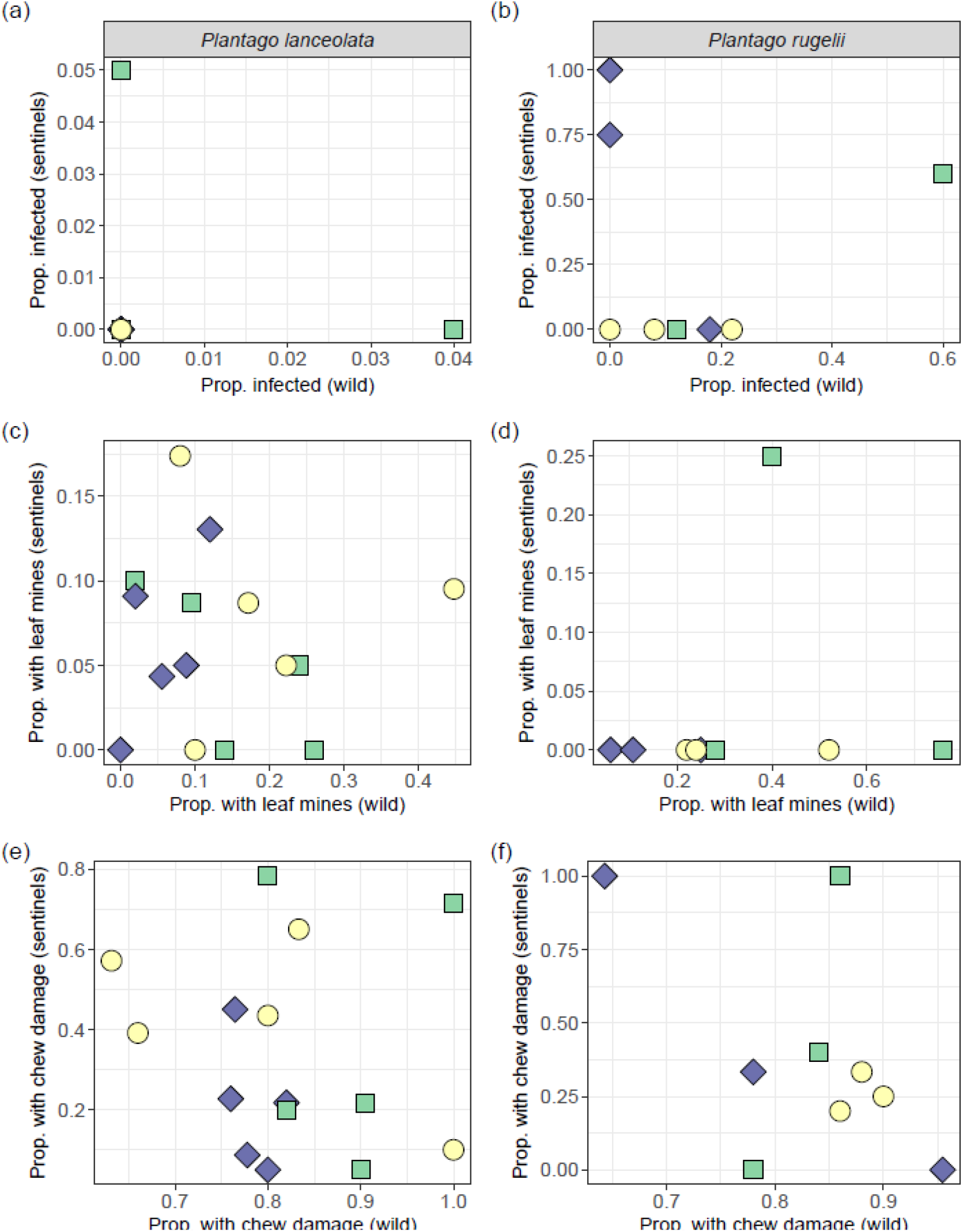
Proportion of sentinels with (**a, b**) powdery mildew infection, (**c, d**) leaf mines, and (**e, f**) chewing damage in relation to the proportion of wild conspecifics with the same type of damage in each site in July 2020. There are 15 sites plotted in each panel, but 12 points with no mildew on either wild or sentinel plants (i.e., overlapping at the origin) in panel (**a**).

Many *P. lanceolata* sentinels experienced herbivory during their week in the field; however, there were no differences between site types for severity (proportion of leaves damaged) of either leaf mines (□^2^ = 4.17, df = 2, P = 0.12) or chewing (□^2^ = 3.82, df = 2, P = 0.15). Prevalences of leaf mines and chewing (proportion of plants affected) were not correlated between sentinels and wild *P. lanceolata* in the same sites during July 2020 (leaf mines: r = − 0.03, P = 0.92, Fig. **6c**; chewing: r = −0.14, P = 0.62, Fig. **6e**).

All 36 *P. rugelii* sentinels survived to the end of the experiment. Eight of these became infected with powdery mildew in urban sites (Clifton Park and Tilles Park), three became infected in a single suburban site (Kirkwood Park) and none became infected in rural sites (Fig. **6b**). Although this trend was consistent with the greater prevalence of infection observed in more urban populations of wild *P. rugelii* in 2020 (Fig. **3d**), there was no significant correlation between prevalence of infection on sentinels and wild plants surveyed in those sites during July (P = 0.95, Fig. **6b**). Instead, the proportion of infected *P. rugelii* sentinels increased with proximity to wild infected plants within the site (P = 0.048, Fig. **S3**).

Only a single *P. rugelii* sentinel acquired a leaf mine during the field exposure. *Plantago rugelii* sentinels experienced chewing-type herbivory in most field sites (Fig. **6f**), and severity of chewing damage did not differ between site types (□^2^ = 1.25, df = 2, P = 0.53). Prevalence of herbivory on *P. rugelii* sentinels was not correlated with that on wild conspecifics in those sites during July 2020 (leaf mines: r = 0.15, P = 0.71, Fig. **6d**; chewing: r = −0.52, P = 0.15, Fig. **6f**).

## Discussion

We found that the timing of key reproductive events and seasonal prevalence of multiple types of foliar damage for wild herbaceous plants changed along an urbanization gradient. The co-occurring focal plant species responded differently in terms of their reproductive phenology. Specifically, urbanization was associated with earlier flowering and more seed production for *P. rugelii*, but less seed maturation for *P. lanceolata*. These species-specific effects on reproductive phenology were broadly consistent between the two years. Another consistent trend between years was that powdery mildew epidemics on *P. rugelii* started earlier and achieved greater infection prevalence in more urban sites consistently between years. Correspondingly, *P. rugelii* sentinels only became infected in suburban and urban sites. However, there was less mildew on *P. lanceolata*, including for sentinels placed into the field sites. There was no significant effect of urbanization on infection prevalence for *P. lanceolata*. Urbanization was associated with accelerated early-summer herbivory on both plant species but was not associated with herbivory of wild or sentinel plants later in summer. Finally, prevalence of mowing damage on leaves was generally greater in urban and suburban than rural sites.

Reproductive development of *P. rugelii* was accelerated along the urbanization gradient. However, there were neutral-to-negative associations with urbanization on reproductive development of *P. lanceolata*. One possible explanation for this difference involves the differential impact of mowing on flower stems of these species. *Plantago rugelii* flowers are produced along most of the length of the flower stem, starting below the height of mower blades. If the upper portion of the flower spike had been removed by mowing, we recorded the maturity of the flowers or seeds that remained. By contrast, *P. lanceolata* produces taller stems with flowers produced only at the tip, where they are highly susceptible to complete removal by lawn mowing. Indeed, stems with completely removed flower spikes were more common for *P. lanceolata* than *P. rugelii* (Fig. **2**). Therefore, more frequent mowing in suburban and urban sites may have reduced the abundance of flower and seed spikes on *P. lanceolata* but not affected the proportion of *P. rugelii* individuals with flowers or seeds. How variation in frequency and/or height of mowing impacts plant reproductive phenology across urbanization gradients is an open question. Given the consistently shorter length of the flowering season of *P. lanceolata* compared to *P. rugelii*, their different responses to urbanization may also be explained by differences in their responses to environmental cues including temperature and light.

Experimental tests of these possible underlying mechanisms are needed to clarify why urbanization has species-specific effects on reproduction. Moreover, for some plants, later flowering and senescence in urban sites is adaptive (Lambrecht et al., 2016). Thus, future experiments testing the genetic basis for differences in phenology between *Plantago* populations will contribute to our understanding of how plant reproductive traits evolve in cities.

Urbanization was associated with earlier and overall larger powdery mildew epidemics on *P. rugelii*. There are several non-mutually exclusive possible reasons for that trend. Two of those possible reasons involve temperature. First, mildew survival is decreased by exposure to freezing air temperatures (Penczykowski et al., 2015). Therefore, the UHI may have increased mildew overwinter survival. Second, warmer temperatures during the growing season can promote mildew growth and transmission. Because all the field sites were parks and nature areas, the magnitude of the UHI experienced by our focal plants might have been tempered by the presence of trees and other vegetation. However, soil temperatures in the pots of sentinel plants increased from rural to suburban to urban sites (Fig. **S4**), consistent with long-term trends in this region (Fig. **1b**). Thus, as for powdery mildew on oaks (van Dijk et al., 2022), warmer temperatures may have contributed to the increased infection prevalence of *P. rugelii* in suburban and urban populations.

Greater rates of spore arrival to more highly connected urban sites may result in increased mildew prevalence with urbanization. Dispersal along roadside populations could allow mildew to spread between sites over the growing season (Numminen & Laine, 2020) with denser urban road networks bringing more mildew spores. However, arriving spores are only successful at infecting if they land on susceptible plant genotypes, and *Plantago* populations can vary in their susceptibility to arriving mildew strains (Höckerstedt et al., 2018; Jousimo et al., 2014; Laine, 2004). Greater resistance of rural *P. rugelii* to mildew could also have decreased their prevalence of infection, even if rates of spore arrival were equally high across the land use gradient. The fact that none of the *P. rugelii* sentinel plants became infected in rural sites is consistent with low spore arrival in those sites, though we note that sample sizes of *P. rugelii* sentinels were small. At the same time, there were lower rates of infection on wild *P. lanceolata* compared to *P. rugelii* growing in the same sites, as well as near-zero prevalence of infection among the >300 *P. lanceolata* sentinels placed across the urbanization gradient. These results suggest species-specificity of the pathogen strains, and lower regional abundance of spores able to infect *P. lanceolata*. Aerial spore sampling and genotyping of mildew infections collected from both *Plantago* species across the urbanization gradient will provide further insights into the connectivity of pathogen populations within and between land use types.

We hypothesized that early-summer prevalence of insect herbivory would be elevated in urban populations, due to UHI accelerating insect emergence and foraging activity in spring. Indeed, both types of herbivory damage were greatest on urban *Plantago* in our earliest survey month (June). However, we further hypothesized that urban sites would continue to experience greater herbivory over the summer, and that herbivory damage on sentinel plants would similarly increase with urbanization. These hypotheses for later-summer patterns are not supported by our data. Instead, prevalence of leaf mines became similar across site types, peaking at a moderate level in July-August. Prevalence of chewing damage was sustained at a high level across site types for the remainder of the growing season. Rates of herbivory on sentinels in July neither followed a trend with urbanization nor correlated with prevalence of herbivory in the wild populations at that time. Generally, our results are consistent with herbarium evidence that increases in winter temperature are a strong driver of increases in insect herbivory (Meineke et al., 2019). Additional experiments will be needed to tease apart effects of winter and spring climate variables, artificial light at night, and other abiotic and biotic facets of urbanization as contemporary drivers of increased early-summer herbivory on *Plantago* hosts.

As urban areas continue to grow in size and prominence worldwide, there is a societal need for understanding effects of urban environmental factors on plants (United Nations, 2018). Urban vegetation provides ecosystem services that mitigate heat, air pollution, and flooding (Heidt & Neef, 2008). Access to green space is also linked to mental and physical health benefits for humans, including decreased heart rate, increased physical activity, and improved mood (Kondo et al., 2018). The abundance, health, and connectivity of native and introduced plant species are also key in supporting diversity of urban fauna (Johnson and Munshi-South, 2017). Thus, it is critical to resolve the mechanisms by which urban environments impact their fitness. In this study, we showed how plant reproductive development and prevalence of disease, herbivory, and mowing damage varied through time across an urbanization gradient. Our findings point to several mechanisms by which changes in land use may impact plant population growth. The generality of these effects should be evaluated through similar studies replicated across additional urban areas.

## Acknowledgements

We thank Whitney Anthonysamy, Beth Biro, and Solny Adalsteinsson for help selecting field sites and Michelle Pollowitz, Armando Sánchez-Conde, Philippa Tanford, and Shayna Rosenbloom for assistance with field work. Carrie Goodson provided greenhouse and lab support. Mike Dyer and Kim Medley provided greenhouse and hoop house support on the Washington University in St. Louis campus and at Tyson Research Center (TRC), respectively. Susan Flowers provided undergraduate mentoring support through TRC. Bill Winston assisted with acquisition of land use data. We thank staff at St. Louis City Department of Parks, Recreation, and Forestry; Forest Park Forever; Tower Grove Park; Webster Groves Parks and Recreation Department; St. Louis County Parks and Recreation; Kirkwood Parks and Recreation Department; Missouri Department of Conservation; TRC; Missouri State Parks; and Missouri Botanical Garden’s Shaw Nature Reserve for access to field sites.

## Author Contributions

QNF, MJB, and RMP conceived of the study design. QNF, MJB, EG, OSS, and KNF performed field surveys. QNF, OSS, and KNF performed the sentinel experiment. RMP provided logistical support for the field surveys and sentinel experiment. QNF and RMP performed statistical analyses. QNF wrote the first draft, and all authors contributed to subsequent drafts of the manuscript.

## Data Availability

Data and code for analyses will be archived at Dryad upon acceptance of the article.

