## Supporting Information for "Phenology of plant reproduction, foliar infection, and herbivory change along an urbanization gradient"

**Table S1.** Site names, locations (in Missouri, USA), management, site type classifications, and survey years in our study.

| **Site ID** | **Site name** | **Longitude** | **Latitude** | **Municipality** | **Managers** | **Site type** | **Years** |
| --- | --- | --- | --- | --- | --- | --- | --- |
| 1 | Shaw Nature Reserve, Entrance | -90.8237 | 38.48183 | Gray Summit | a | Rural | 2019-20 |
| 2 | Shaw Nature Reserve, Interior | -90.8222 | 38.46734 | Gray Summit | a | Rural | 2019-20 |
| 3 | Route 66 State Park | -90.6037 | 38.5044 | Eureka | b | Rural | 2019-20 |
| 4 | West Tyson Park, Chubb Trail | -90.5868 | 38.52826 | Eureka | c | Rural | 2019-20 |
| 5 | Tyson Research Center, 2, East | -90.5582 | 38.52468 | Eureka | d | Rural | 2019-20 |
| 6 | Tyson Research Center, 1, West | -90.5621 | 38.52729 | Eureka | d | Rural | 2019-20 |
| 7 | Buder South Park | -90.4888 | 38.53711 | Valley Park | c | Suburban | 2019-20 |
| 8 | Simpson Park | -90.4683 | 38.55491 | Valley Park | c | Suburban | 2019-20 |
| 9 | Emmenegger Nature Park | -90.433 | 38.54547 | Kirkwood | c | Suburban | 2019-20 |
| 10 | Powder Valley Conservation Nature Center | -90.4282 | 38.55563 | Kirkwood | e | Suburban | 2019-20 |
| 11 | Kirkwood Park | -90.4207 | 38.58161 | Kirkwood | f | Suburban | 2019-20 |
| 12 | Tilles Park, Ladue | -90.3678 | 38.62151 | Ladue | c | Suburban | 2019-20 |
| 13 | Southwest Park | -90.3676 | 38.57288 | Webster Groves | g | Suburban | 2019-20 |
| 14 | Blackburn Park | -90.3418 | 38.58253 | Webster Groves | g | Suburban | 2019-20 |
| 15 | Francis Park | -90.3037 | 38.58729 | St. Louis City | h | Urban | 2019-20 |
| 16 | Clifton Heights Park | -90.2915 | 38.61437 | St. Louis City | h | Urban | 2019-20 |
| 17 | Tilles Park | -90.2906 | 38.60139 | St. Louis City | h | Urban | 2019-20 |
| 18 | Forest Park | -90.2708 | 38.63841 | St. Louis City | h,i | Urban | 2020 |
| 19 | Tower Grove Park, 2, West | -90.2667 | 38.60833 | St. Louis City | h,j | Urban | 2019-20 |
| 20 | Tower Grove Park, 1, East | -90.2505 | 38.60567 | St. Louis City | h,j | Urban | 2019-20 |
| 21 | Benton Park | -90.2242 | 38.5972 | St. Louis City | h | Urban | 2019-20 |
| 22 | Lafayette Park | -90.2176 | 38.61526 | St. Louis City | h | Urban | 2019-20 |

**Site managers:** a. Missouri Botanical Garden; b. Missouri State Parks, a division of the Missouri Department of Natural Resources; c. St. Louis County Parks and Recreation; d. Tyson Research Center, Washington University in St. Louis; e. Missouri Department of Conservation; f. Kirkwood Parks and Recreation Department; g. Webster Groves Parks and Recreation Department; h. St. Louis City Department of Parks, Recreation, and Forestry; i. Forest Park Forever; j. Tower Grove Park

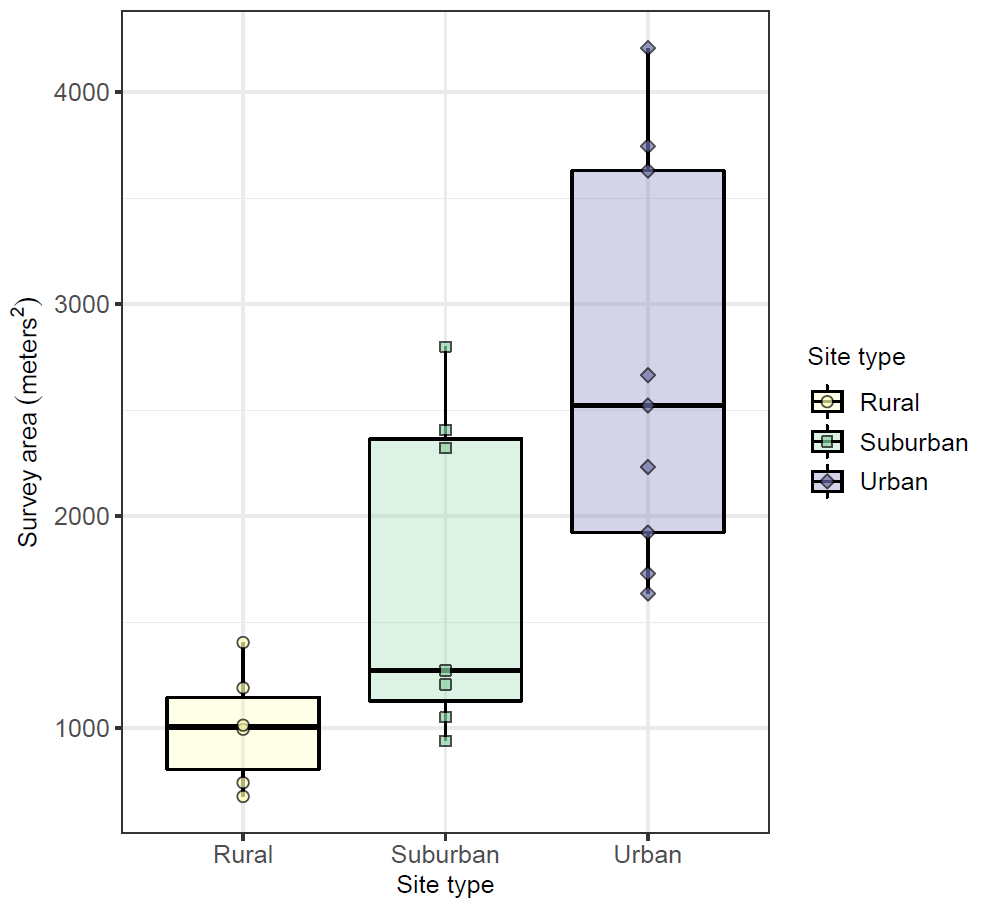

**Figure S1.** Total area (m^2^) surveyed in each field site.

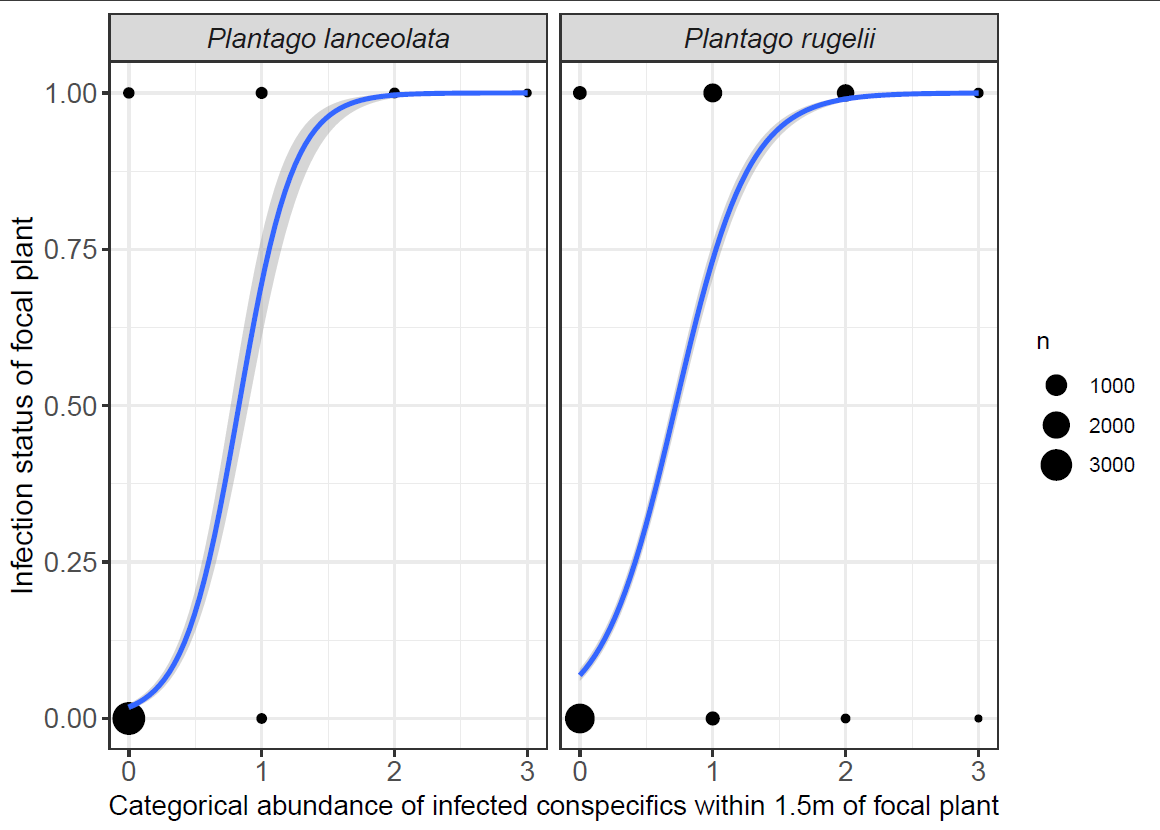

**Figure S2.** Binary infection status of focal surveyed plant (y-axis) vs. categorical abundance of infected conspecifics within a 1.5 m radius (x-axis categories: "0" = 0, “1” = 1-10, “2” = 11-50, and “3” = 51-100 infected conspecifics). The plotted data are from all field sites and survey dates, beginning in July 2020. Size of black circles represents the total number of focal plants (n) with overlapping data points.

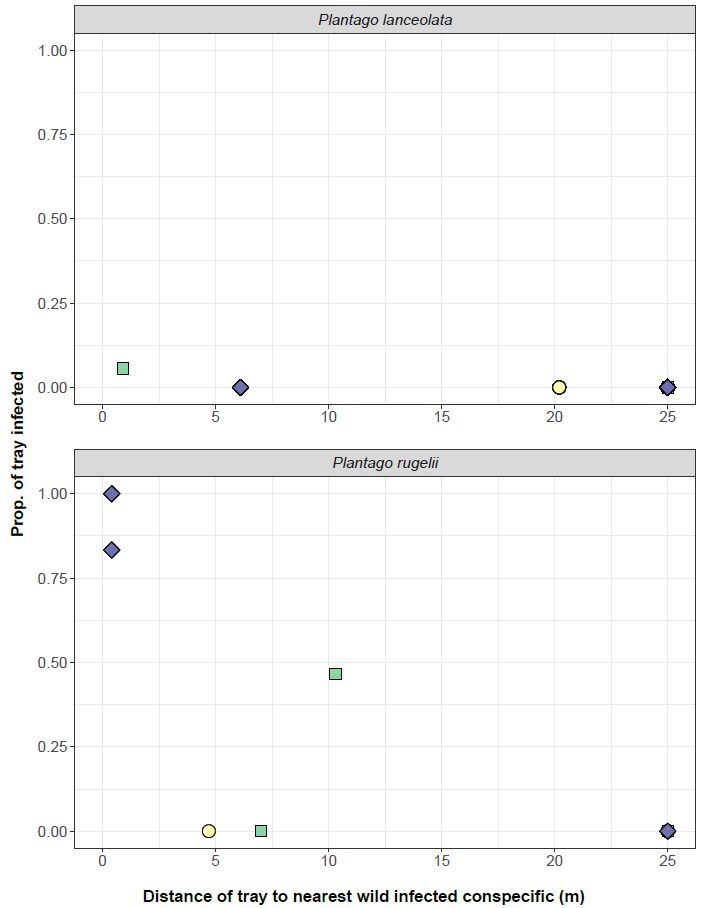

**Figure S3.** Distance (m) from trays of sentinel plants to the nearest infected conspecific in each field site in July 2020. The radius of our search for mildew extended to 25 m from the trays of sentinels. In multiple sites, there were no wild infected plants within that search radius. We assigned those sites the maximum value of 25 m for the purposes of this plot and accompanying statistical analyses. Thus, the overlapping points at x = 25 m include sites where the actual distance to infected plants may have been much greater.

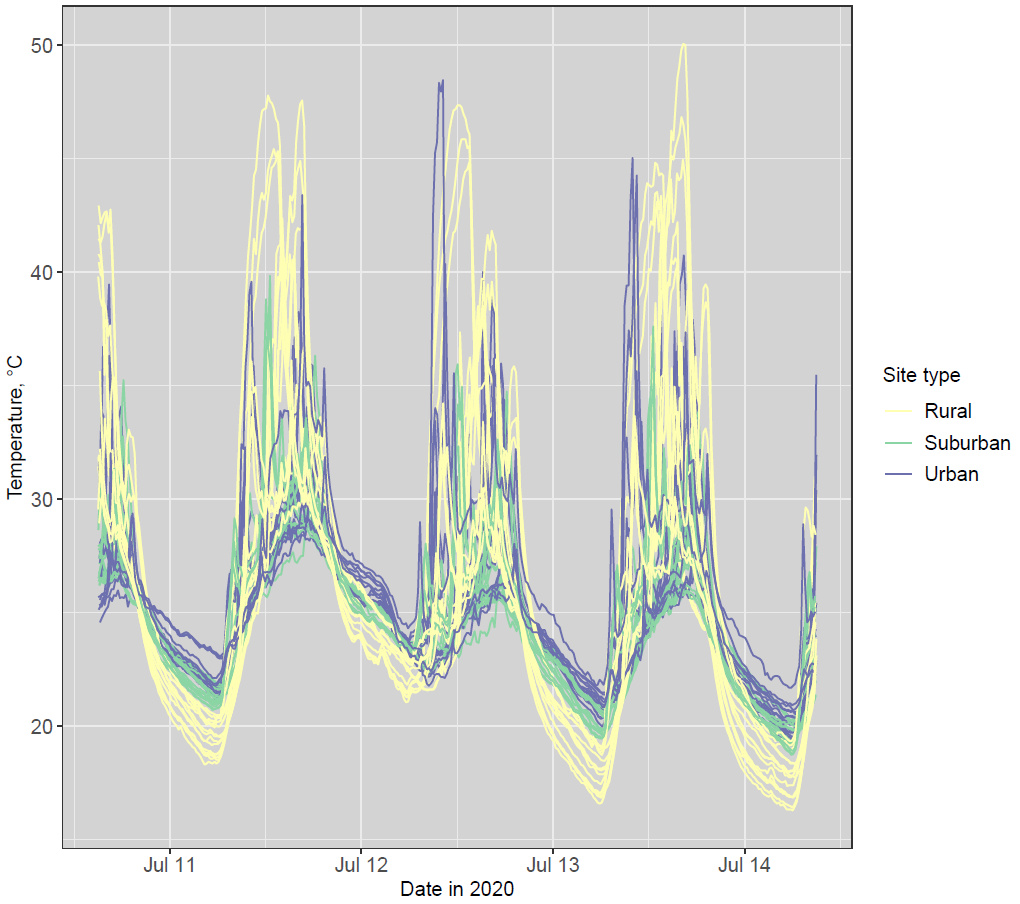

**Figure S4.** Temperature (°C) readings of HOBO MX2201 dataloggers in pots of sentinel plants within rural (yellow), suburban (green), and urban (purple) sites. Vertical grid lines at date label tick marks indicate midnight (00:00), and vertical lines between date labels indicate noon (12:00). Dataloggers were initially placed 1 cm below the soil surface in each pot; however, exceptionally high and inconsistent daytime temperature readings suggest that the dataloggers became exposed to direct sunlight. As such, daytime temperatures should not be interpreted. During nights, when readings were not impacted by direct sunlight, there is a clear pattern of increased nighttime temperature in more urban sites (consistent with the greater average monthly temperature in more urban sites shown in main text Fig. 1b).
